# Colicin-mediated transport of DNA through the iron transporter FepA

**DOI:** 10.1101/2021.05.11.443673

**Authors:** Ruth Cohen-Khait, Ameya Harmalkar, Phuong Pham, Melissa N. Webby, Nicholas G. Housden, Emma Elliston, Jonathan TS. Hopper, Shabaz Mohammed, Carol V. Robinson, Jeffrey J. Gray, Colin Kleanthous

## Abstract

Colicins are protein antibiotics used by bacteria to eliminate competing Escherichia coli. Colicins frequently exploit outer membrane (OM) nutrient transporters to penetrate through the strictly impermeable bacterial cellular envelope. Here, applying live-cell fluorescence imaging we were able to follow colicin B (ColB) into E. coli and localize it within the periplasm. We further demonstrate that single-stranded DNA coupled to ColB is also transported into the periplasm, emphasizing that the import routes of colicins can be exploited to carry large cargo molecules into bacteria. Moreover, we characterize the molecular mechanism of ColB association with its OM receptor FepA, applying a combination of photo-activated crosslinking, mass spectrometry, and structural modeling. We demonstrate that complex formation is coincident with a large-scale conformational change in the colicin. Finally In vivo crosslinking experiments and supplementary simulations of the translocation process indicate that part of the colicin engages active transport by disguising itself to part of the cellular receptor.

---

Bacteria are the most common and diverse form of life on earth. The remarkable abundance of different bacterial strains and species capable of surviving at almost any environment frequently leads to severe inter-bacterial competition over space and limited resources 1. The recurrent need to compete for basic life necessities has led the evolution of efficient nutrient delivery strategies as well as inter-bacterial weapons. Excessive competition for soluble iron ions has led the evolution of the potent iron chelating agents termed Siderophores 2. Most commonly used bacterial weapons are usually toxins which either perform a nuclease activity 3 or destruct the cellular membrane integrity 4. The most challenging step in this antibiotic bacterial war is to specifically deliver the toxin to the closely related competing bacterial strain while avoiding self-killing. Immunity proteins expressed alongside the toxin inactivate the cytotoxic activity within the producing strains 5. Cytotoxic proteins can be delivered either in a contact-dependent manner, targeting neighboring cells relying on the assembly of supramolecular secretion machineries 6, or in a way which does not depend on physical contact between the cells as exemplified by bacteriocins 7.

Colicins are bacteriocins that are specifically active against E. coli. The key obstacle colicins face prior to executing their killing action is penetrating the impermeable bacterial cellular envelope 8. Other antibiotic molecules as well as Siderophores and other large nutrient molecules face a similar challenge. The cellular envelope of gram negative bacteria is composed of the non-energized outer-membrane (OM), the energized phospholipidic inner-membrane (IM) and the intervening peptidoglycan containing periplasm 9. Moreover, colicins are large (29-75 kDa) proteins that cannot spontaneously diffuse through the cellular envelope 10, hence they also face the challenge of finding active transport into the cell, foremostly the OM 11. There are two structurally related energy transfer systems that are exploited by colicins to traverse the OM barrier: 1) The Tol system, which stabilizes the OM during cellular division 12, and 2) the Ton system composed of the IM protein complex TonB-ExbB-ExbD, which is mainly responsible for the active import of nutrients such as siderophores through specialized OM receptors 13.

Most of a colicin’s structure is involved in overcoming the translocation impediments across the cellular envelope. Colicin proteins are usually composed of three major structural domains: 1) a central receptor binding domain (R) 2) an N-terminal translocation domain (T) and 3) a C-terminal cytotoxic domain 14. The main purpose of the receptor binding domain is to interact with a specific receptor (usually a nutrient transporter) in the OM and to anchor the colicin to the cell. Once the colicin is in position the T domain interacts with a different OM protein, upon which the colicin translocation is dependent 15, 16. Colicin B (ColB) is a pore forming toxin that is one of the earliest colicins to be described 17. However, little is known about the cellular translocation process of ColB beyond its dependence on the OM ferric enterobactin transporter FepA, and the Ton system 18. No additional proteins have been identified for ColB toxicity, which may explain why, unlike most other colicins, ColB is composed of only two functional domains: an N-terminal domain that serves as both a receptor binding domain and a translocation domain (ColB RT) and a pore forming C-terminal cytotoxic domain 19. The ColB receptor FepA is a 22-stranded β-barrel TonB-dependent transporter (TBDT) with an N-terminal plug domain blocking its lumen 20 21, 22.

Here, we examined whether ColB could be exploited for delivery of non-proteinaceous cargo in the form of ssDNA into bacterial cells. We have then examined the molecular mechanism of the ColB interaction with its receptor FepA, and the way ColB exploits FepA to engage active transport into the cell. Applying live-cell fluorescence microscopy we exhibit the direct visualization of translocated ColB RT domain in the periplasm of E. coli, demonstrating unambiguously that translocation requires only FepA and TonB. Furthermore we show that the active transport mechanism for ColB through FepA can be exploited to deliver ssDNA into the cell. We then applied a unique approach combining in vitro and in vivo photoactivated crosslinking, mass-spectrometry, and structural modeling with Rosetta to follow the key stages in the ColB-FepA association process. We identify an energetically favorable encounter complex that induces major conformational changes in the colicin and consequently leads to the formation of a stable complex in vitro. Following further in vivo crosslinking data with additional structural modeling, we identify an intermediate TonB dependent translocation state. We show how the partially unstructured N-terminal 55 amino acids of the colicin replace the N-terminal half plug domain of the receptor, which is actively unfolded by TonB. This ultimately allows direct engagement of the colicin with the periplasmic energy transferring agent, which emphasize how this system is suitable for conjugated cargo transport.

## ColB can transport ssDNA through FepA

We examined whether the ColB RT domain could be followed into bacterial cells. We have conjugated the ColB RT domain (residues 1-341) to a small fluorescent dye (Alexa 488 nm) and applied live-cell fluorescence microscopy to examine its ability to penetrate into E. coli cells, as well as determine its cellular localization (Fig 1A). We defined cellular penetration as a fluorescent signal resistant to trypsin treatment. Deletion of the ColB TonB box (residues 17-21) resulted in a complete loss of the colicin’s ability to translocate, however it has maintained its ability to bind FepA (Fig 1A). No binding was detected in FepA knock out cells (Fig 1A). Consistent with ColB being a pore forming colicin, the fluorescent signal was lost upon cellular spheroplasting (disruption of the OM and the periplasmic peptidoglycan layer) suggesting that ColB RT remained in the periplasm and did not translocate into the cytoplasm (Fig 1A). In order to examine whether colicins could be used to deliver macromolecules into the cell in a similar way to siderophores 23 we generated fluorescent colicin-DNA fusions. We used maleimide conjugation to attach fluorescent ssDNA (15A or 5A 10C) to an introduced C-terminal Cys on ColB RT. We verified that the fluorescent signal followed the attached DNA molecule by exposing it to DNase treatment (Supplementary Fig 5). Hence, any translocated fluorescent signal indicated that the conjugated DNA molecule has transferred into the cell. The fluorescent signal appeared to follow the ColB RT periplasmic localization in a FepA dependent manner (Fig 1B). In order to test whether dsDNA could follow the colicin’s path into the cell the ColB RT 5A 10C construct was incubated with a 10G DNA fragment. The addition of the 10G fragment did not disrupt the construct’s ability to translocate (Fig 1B). However it is not clear whether the dsDNA has fully translocated into the cell, or whether the applied force (due to TonB activity) has disrupted the dsDNA - so that only the conjugated 5A 10C fragment has translocated in.

**Fig. 1.**
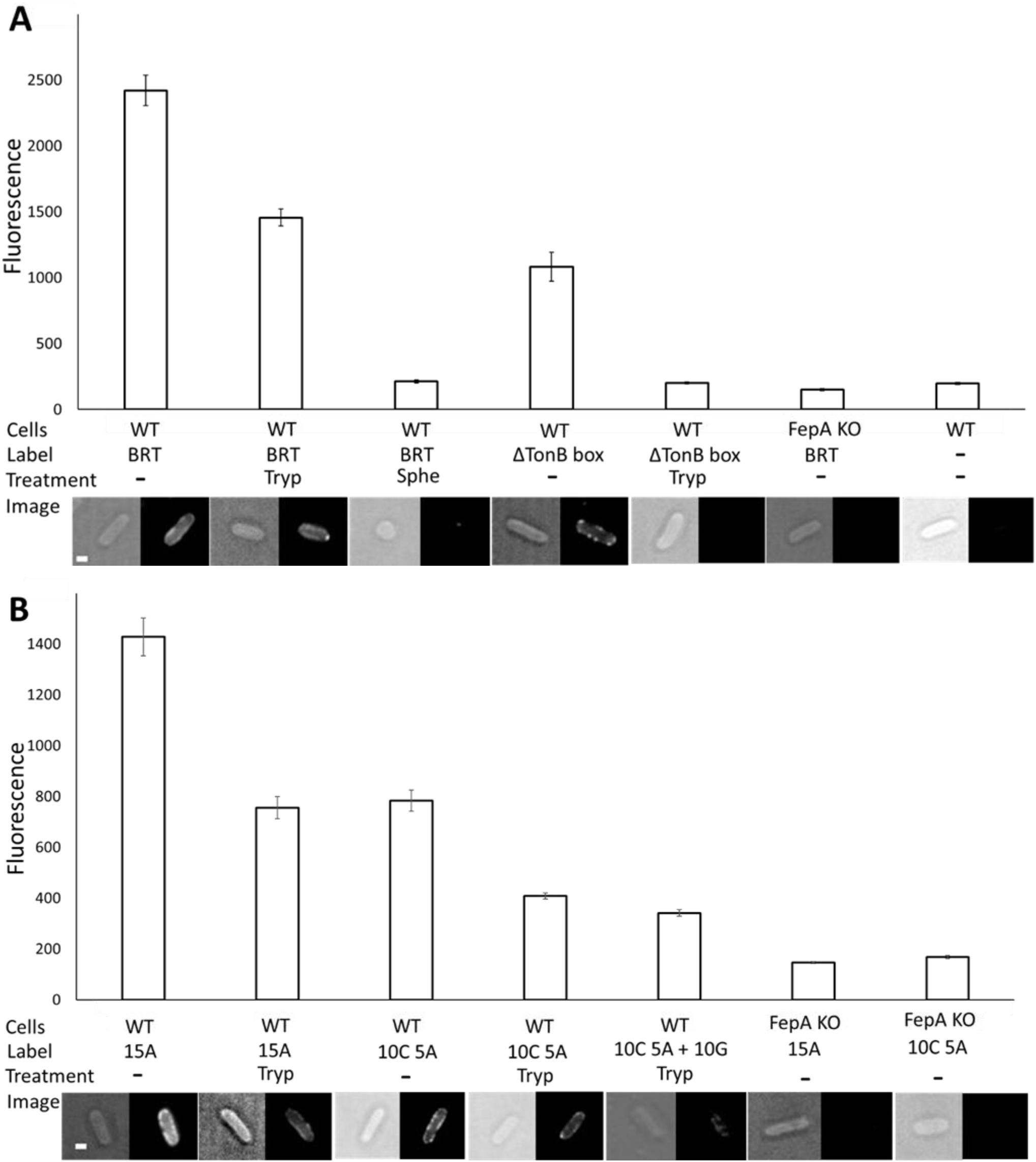
ssDNA follows the ColB RT translocation path into *E. coli* cells. **A.** Translocation of ColB RT-Alexa 488 (BRT) or ColB RT ΔTonB box – Alexa 488 (ΔTonB box) constructs into *E. coli* MC1655 cells (WT) or *E. coli* BW25113 ΔFepA (FepA KO) cells grown in minimal media to mid log growth phase (OD_600_~0.35). Cellular translocation defined as fluorescent signal resistant to trypsin treatment (Tryp). Cytoplasmic localization defined as fluorescent signal remaining after spheroplasting the cells, which results in the removal of the OM and the periplasmic peptidoglycan layer (Sphe). The averaged fluorescence intensities were calculated from at least 120 cells (30 cells x 4 biological repeats), standard error bars of each treatment are shown. Representative cellular images below each treatment. Scale bar - 1 μm. **B.** Translocation of ColB RT-DNA fused constructs: ColB RT-15A Alexa 488 (15A), ColB RT 10C 5A Alexa 488 (10C 5A), ColB RT 10C 5A Alexa 488 + 10G (10C 5A + 10G).

## Receptor binding induces large-scale conformational changes in ColB

In order to examine the stoichiometric complex ratio of the FepA-ColB complex we applied advanced native state mass-spectrometry on a complex assembled in vivo on E. coli surface. The FepA-ColB complex ratio was 1:1 (Supplementary Figure 1) suggesting that ColB associates and translocates via a single copy of FepA. We have further applied Rosetta based structural simulations to determine how the ColB RT domain associates with FepA. We examined the available PDB structures of ColB 24 (PDB 1RH1) and FepA 20 (PDB 1FEP) applying a Rosetta docking algorithm 25, which revealed a clear energy funnel (Fig 2D) for an encounter complex (EC) structure (Fig 2A). We were able to back up the Rosetta model predictions applying a pBPA crosslinking approach. We introduced P-benzoyl-L-phenylalanine (pBPA) mutations on unique ColB RT surface loops, previously highlighted as potential FepA binding sites 26 (Supplementary Fig 2). Exposure to UV (365 nm) results in pBPA non-specific crosslinking to a carbon atom within ~4 Å 27. We executed photoactivated crosslinking experiments both in vitro using the OM protein fraction as a FepA source, or in vivo on live E. coli cells. We identified crosslinks on SDS-gels and further analyzed them by LC MS-MS, as previously described in White et al 28. Applying this method we identified 3 crosslinks in vitro – two of which (ColB 202 & 205 with FepA 642 & 639 respectively) supported the initial EC computed with Rosetta (Fig 2A, Supplementary Fig 3A, Supplementary Fig 4A-C). Thus, recent progress in the Rosetta docking energy function 25, 29 allowed the accurate prediction of the EC (Fig 2A) as has been further confirmed by pBPA crosslinking experiments (Fig 1, Supplementary Fig 3A, Supplementary Fig 4A-C). However the third crosslink (ColB 55 with FepA 652) yet remained to be explained.

**Fig. 2.**
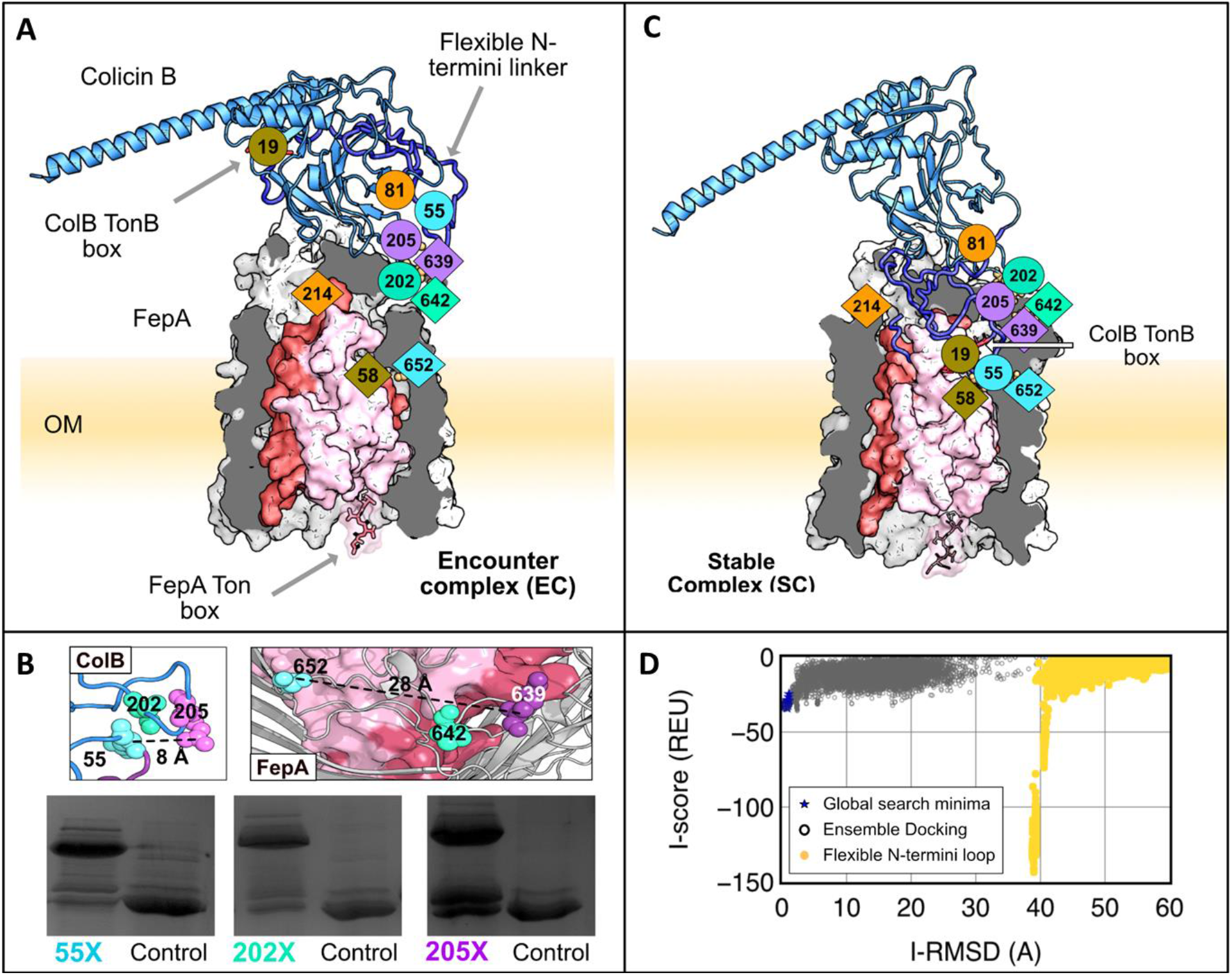
Structural insights on the ColB-FepA complex by pBPA crosslinking and Rosetta-based structural modeling. **A.** Initial encounter complex (EC) modeled with moderate to little backbone flexibility (under 5 Å RMSD). ColB (blue) and FepA (grey) form this encounter complex with *in vitro* crosslinks, FepA-K639 and ColB-D202 (teal), and FepA-P642 and ColB-R205 (purple), which lie in proximity in the model. The last *in vitro* crosslink pair, FepA-S652 and ColB-Q55 (cyan), and the two *in vivo* crosslinks, FepA-T58 and ColB-M19 (olive), and FepA A214 and ColB-G81 (orange), are not satisfied in this structure. **B.** Mapped *in vitro* crosslinking sites on the ColB and FepA PDB structures (1RH1 and 1FEP respectively). Cropped relevant crosslinks gels. Self-crosslinking control to the right of each lane (full *in vitro* crosslinking image at Supplementary Fig 3A). **C.** Fully assembled spontaneously formed stable complex (SC) modeled with the Rosetta FlopyTail algorithm^58^ stimulating the partially unstructured ColB 1-55 as a floppy-tail. **D.** Rosetta Interface score vs Interface RMSD for output structures identified by local docking (ReplicaDock2) of ColB to FepA. RMSD is measured relative to the lowest-scoring global docking structure. There is a deep minima resulting from the arrangement of the flexible N-linker for the FloppyTail models.

ColB 55 appears in close proximity to ColB 202 & 205 on the ColB PDB structure, yet our crosslinking data show that it does not crosslink to neighboring residues on the receptor – suggesting that complex formation induces conformational changes within the colicin (Fig 2B). In order to improve the structural model of the ColB-FepA complex, we stimulated the partially unstructured N-terminal tail of ColB (residues 1-55) as a ‘floppy tail’, allowing it to sample its environment freely 25. The resulting model of the stable complex (SC) explains all three in vitro observed crosslinks (Fig. 1C) and seems to be energetically favorable over the initially calculated EC (Fig 2D, Movie 1). The calculated SC seems to also bring the ColB TonB box closer to the FepA lumen (Fig 2C).

## ColB exploits FepA for its active translocation into the cell

The route of ColB translocation in a FepA dependent manner is unknown. Here we show how the partially unstructured flexible N-terminal tail of ColB (residues 1-55) occupies the TonB dependent channel generated by the FepA half plug domain unfolding (Fig 3). We suggest that while complex formation (Fig 2) is a highly specific step, the translocation mechanism through 22 stranded beta-barel TBDTs is likely to be applicable to many other systems sharing similar protein folds (Supplementary Fig 6). We identified the three crosslinks observed in vitro also in vivo as well as additional two crosslinks which we further mapped by LC MS-MS (Supplementary Fig 3B, Supplementary Fig 4D-E). The additional two crosslinks did not form in the absence of the energy transferring agent TonB (Supplementary Fig 3C).

**Fig. 3.**
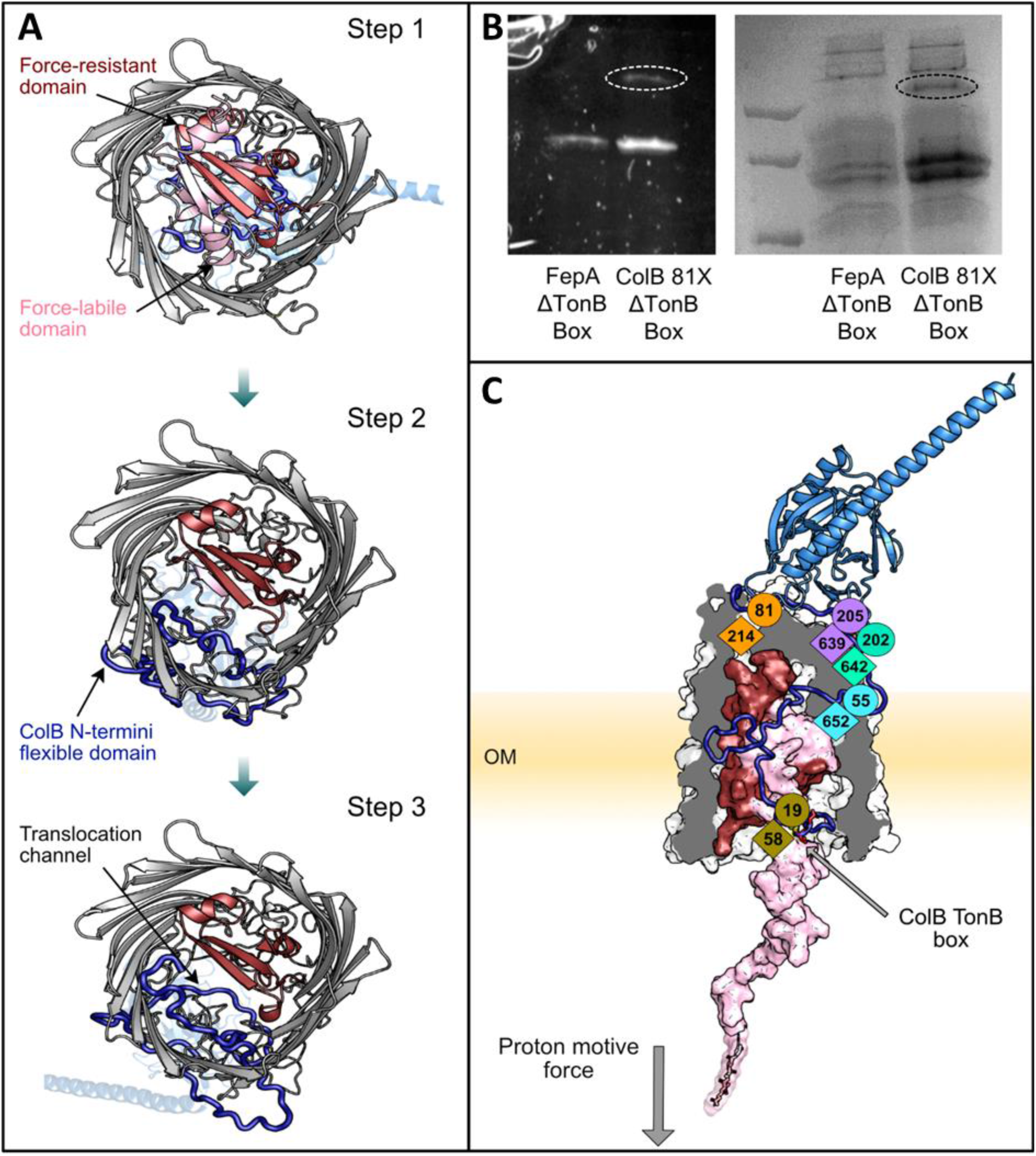
The partially unstructured N-terminal ColB RT tail occupies the gap generated by the active unfolding of the FepA N-terminal half plug domain. **A**. A bottom-to-top view of the hypothesized translocation pathway (Stage 3) created with Rosetta by pulling the FepA N-terminus into the cell. Step 1: SC complex is formed and the force-labile half-plug domain (light pink) begins to unfold. Step 2: The force-labile half-plug is partially unfolded which allows the ColB N-terminal loop (blue) to occupy the void created by the absence of the plug domain. Step 3: The unfolding of the FepA half-plug domain creates a channel for the ColB N-terminal loop to enter. **B**. The ability of ColB-81X GFP to crosslink *in vivo* as a function of both ColB and FepA TonB boxes. GFP fluorescence (right) Coomassie blue stain (left). Crosslinked band circled. **C**. The top-scoring model portraying the translocon state (Step 3). In this final-stage model, all the crosslink constraints are satisfied and the model is energetically favorable over other states.

The TonB box is a conserved penta-peptide sequence essential for interaction with TonB 30, and thus is expected to lead the colicin translocation course. Two essential TonB boxes participate in the ColB translocation process: one on the colicin itself and the other on its OM receptor FepA 18,31. We examined the ability of the in vivo observed ColB 81-FepA 214 crosslink to form as a function of both the FepA and ColB TonB boxes. The ColB 81-FepA 214 does not form in the absence of the FepA TonB box, but it still forms in the absence of the ColB TonB-box (Fig 3B). Hence, as both TonB boxes are essential for full colicin translocation, the ColB 81-FepA 214 crosslink appear to capture a stable intermediate translocation step. These experiments were not performed on the second in vivo identified crosslink ColB 19-FepA 58 as ColB 19 is already part of the ColB TonB box.

In order to investigate the structures during the dynamic translocation process, we applied Rosetta to stimulate the unfolding of the N-terminal half (residues 1-74) of the globular FepA plug domain as previously demonstrated on BtuB32. We stimulated the ColB-FepA translocation process starting with the computed SC structure (Fig 2C) and using the in vivo identified crosslinks as guides to generate three intermediate structures in 4 Å increments (Fig 3A, Fig 3C). The stimulated structures indicate that the translocating N-terminal ColB tail (residues 1-55) occupies the cavity generated by the FepA half plug removal with the ColB TonB box now positioned in place of the former FepA TonB box (Fig 3A, Fig 3C).

## Discussion

The OM is the foremost barrier into the cell which due to its low permeability provides gram negative bacteria with protection against a various range of antibiotics 33. A recently developed antibiotic delivery strategy termed ‘The Trojan Horse’ relies on antibiotic molecules conjugation to siderophores which are actively consumed by bacteria due to bacterial innate need for iron ions 23,34. Here we addressed the question of whether ColB could be exploited to deliver non-proteinaceous molecules into bacteria. Firstly we applied live-cell fluorescent microscopy to visualize ColB RT in the periplasm of E.coli (Fig 1A). We were then able to identify fluorescent ssDNA molecules covalently attached to ColB RT in the periplasm (Fig 1B). We show that the ssDNA translocation process depends on FepA in a similar manner to the translocation process of ColB (Fig 1B).

The FepA-ColB interaction occurs via two common protein folds: FepA like many other OM transporters is a 22 stranded β-barrel with a globular N-terminal plug domain 35. ColB RT shares 96% sequence similarity with the N-terminal domain of colicin D (ColD) and thus is believed to take a similar route into the cell. However as ColD is a tRNase its cytotoxic domain needs to further be delivered into the cytoplasm while the periplasm is the destination of the pore forming ColB. ColB RT belongs to the pyocin_S domain superfamily 36, a 3D protein fold that is very common amongst bacteriocins with nuclease cytotoxic activity (such as ColD) and is usually associated with translocation across the IM. ColB is the only known example of a pore forming colicin exhibiting the pyocin_S domain, which apparently uses this fold to translocate across the OM. The surprisingly high structural similarities these proteins share with other OM transporters/colicins is emphasized in Supplementary Fig 6. Nevertheless the FepA-ColB complex is highly specific. While ColB exclusively interacts with FepA (Fig 1A), other pyocin_S domains do not (Cao & Klebba, 2002; Wookey & Rosenberg, 1978) 37, 38.

ColB depends on TonB for its cellular penetration, yet it is not clear how ColB hitchhikes the TonB system to promote its active transport into the cell. It has been proposed that ColB passes through the FepA lumen 18, this hypothesis has been supported by the partial dislocation of the FepA half plug domain demonstrated by exposure to periplasmic Cys labelling 18, 39. On the other hand when a different Cys label was used it appeared that ColB binding induced no conformational changes in the FepA plug domain, implying that ColB probably does not penetrate the cell through FepA 40.

The TonB box is a conserved penta-peptide sequence essential for interaction with the TonB energy transferring system (Schramm et al, 1987) and thus is expected to lead the colicin translocation process. Two essential TonB boxes participate in the ColB translocation process: one on the colicin itself and the other on its OM receptor FepA (Devanathan & Postle, 2007; Mende & Braun, 1990). Here we showed that while the FepA TonB box is essential for the in vivo observed crosslink of ColB 81 with FepA 214, the ColB TonB box is not (Fig 3B, Supplementary Fig 3B Supplementary Fig 4E), implying that the colicin is stuck in a stable intermediate translocation step.

It has been shown on the FepA homologous vitamin B12 transporter BtuB, that the N-terminal globular plug domain of the receptor is composed of two mechanically independent half-plug domains, with the N-terminal half plug more likely to unfold due to TonB applied force 32. We have simulated the unfolding of the N-terminal plug domain of FepA as previously emphasized on BtuB applying AFM (Hickman et al, 2017). The computed model (Fig 2C) emphasizes the importance of the two independent encounters with the energy transferring agent TonB: The first receptor mediated encounter allows the translocation of the ColB TonB box into the periplasm (Fig 3C), and later a second encounter allows the active colicin translocation into the cell. The computed model demonstrates how the N-terminal tail of the translocating colicin mimics the unfolded receptor half-plug domain and replaces the receptor’s TonB box with the colicin’s one (Fig 3A, Fig 3C).

Previous studies have emphasized the importance of the FepA surface loops in substrate recognition and have further indicated the sequential order of the loops movement upon substrate binding 41–43. Here we did not account for possible energetically favorable movements of the FepA surface loops because our experimental data focused on the binding colicin, and the manner it exploits FepA for active transport into the cell. Moreover, the mapped crosslinking area on FepA has not been previously identified as vital for the ColB-FepA interaction. It is possible that ColB binding induces movement in the surface loops of FepA in a similar manner to ferric enterobactin, which may slightly alter the final in vitro SC structure (Fig. 1C). However, it is unlikely to change our main structural conclusions regarding the initial FepA-ColB recognition step EC (Fig 1A), the induced conformational changes in ColB (Fig 2B, Fig 2C) and the essence of the proposed first translocation step across the FepA lumen (Fig 3A).

Overall we show that the translocation process of ColB is a dynamic process that involves receptor mimicry and relies on two energy transfer events. The approach described here, combining photoactivated crosslinking and structural modeling enabled a detailed characterization of the cascade of complex readjustment events that drive colicin transport across the OM. The described tranlocation mechanism likely applies to other FepA and TonB dependent toxins such as ColD and phage H8 44. Moreover, the defined mechanism is also relevant to other bacteriocins utilizing TBDTs with similar folds. The emphasized ability of bacteriocin-DNA conjugations to follow the colicin route into the cell opens a large range of opportunities to utilize bacteriocins for bypassing the otherwise strictly impermeable gram negative bacterial OM. This includes development of novel antibiotics delivery strategies in a similar way to siderophores (of which colicins are natural mimics). It also applies to the area of genomic manipulations in non-domesticated bacteria, as the colicin-DNA fusions provide a highly specific manner of targeted DNA delivery into bacteria.

## Materials and Methods

### Protein expression and purification

All colicin constructs were conjugated to a 6xHis tail at their C-terminus and cloned at the second multiple cloning site of the apACYCDuet-1 (Novagen) plasmid where they were expressed under a T7 promotor. The plasmids were transformed into BL21(DE3) *E. coli* cells. Transformed cells were grown at 37° C in Lysogeny Broth (LB) media pH 7.2 while shaking at 180 rpm to OD_600_~0.6 at which point 1 mM Isopropyl-D-thiogalactopyranoside (IPTG) was added and the temperature was reduced to 20° C for an overnight incubation. The protein has been eluted as described in Khait and Schreiber (2012)^47^. Briefly, protein expressing cells were re-suspended in 20 mM Tris pH 7.5, 0.5 M NaCl, 5 mM Imidazole and sonicated (Sonicator 4000: 70%, 1.5 min, 3 sec on:7 sec off). The sonicated cellular extract was spun down and the supernatant has been incubated with His-binding resin (Merck 69670-5) for 10-30 min at room temperature. The Ni-resin and the bound protein were then gently (1000 g <) spun down, washed three times and re-suspended in the same buffer containing 0.5 M Imidazole which allowed protein elution. The protein has been dialyzed to PBS at 4° C overnight. FepA has been expressed on a pBAD/Myc-HisB (Novagen) plasmid, transformed into either Bl21(DE3) or BW25113 ΔFepA (JW5086-3) *E. coli* cells. FepA has been expressed similarly to the colicin proteins except of the LB growing media pH being 6.12 and protein expression induction with 0.15% L-Arabinose. The FepA containing protein OM fraction has been purified as previously described for ompF^48^. Protein concentrations were determined through absorbance at 280 nm using a sequencebased extinction coefficient.

### Fluorescent protein labelling

Colicins were conjugated to Alexa 488 or 15 b DNA – Alexa 488 by maleimide reactions as described in Kleanthous et al^49^ with some adaptions: the purified protein was incubated with 10 mM DTT for one hour at room temperature (or overnight at 4° C) it was then run throw a desalting column (buffer: 25 mM Tris pH 7.5, 100 mM NaCl) and immediately incubated with x1.1 or x3 ratio of Eurogentech maleimide DNA conjugates or maleimide Alexa 488 respectively for one hour at room temperature, the reaction was terminated by the addition of 5 mM DTT. The protein has been desalted again, retrieved by Ni-beads as in the previous section and dialyzed to PBS. The efficiency of the fluorescent conjugations has been determined on a fluorimeter (UV/VIS V-550 Jasco). The protein-DNA conjugation sensitivity to DNase and trypsin treatments has been analyzed on 15%-SDS page gels.

### Native State Electrospray Ionization Mass-spectrometry

Sixty mg of ColB RT (341 aa) were added to a 5 L culture of BW25113(ΔFepA) cells over expressing FepA from a pBAD/Myc-HisB (Novagen) plasmid. The complex has been purified following the protocol previously described for ompF^48^. A 5 ml HiTrap desalting column (GE Healthcare) was used to exchange the complex buffer into 100 mM ammonium acetate, 1% (w/v) β-OG, pH 6.9. Mass spectrometry measurements were made from a static nanospray emitter, using gold-coated capillaries prepared in-house^50^, on a quadrupole time-of-flight mass spectrometer (Micromass) modified for high mass transmission. Liberation of the protein complex from β-OG detergent required energetic instrument parameters and the low m/z region of spectra where dominated by detergent clusters. Operating conditions used include capillary voltage 1800V, sample cone 200V, extractor 10V, collision cell energy 140-200V, source backing pressure 5.92 x10-3 mbar and Argon collision cell pressure 3.5 – 5 MPa.

### Crosslinking

The crosslinking procedure was similar to White et al (2017)^33^. In short *pBPA* mutations were introduced at 21 different positions of ColB RT (341 aa) GFP. For *in vitro* crosslinking 1 μM of *pBPA* containing colicin was incubated with 1 ml of an OM protein fraction (in PBS pH=6.5, 5 mM EDTA, 2% β-OG) extracted from BW25113 FepA knock-out cells over expressing FepA containing ~1 μM FepA and exposed to UV light (365 nm) for 1 hour at 4¤ C. The colicin and bound/crosslinked FepA were then extracted by EDTA resistant Ni-beads cOmplete (Merk). For *in vivo* crosslinking the colicin was incubated with 800 ml cells over expressing FepA overnight at 20° C. The *pBPA* containing colicin was added to the LB media (pH 6.12) and incubated for 90 min at 37¤ C while shaking. The cells were then spun down, colicin excess was washed with 50 ml PBS, the cells were re-suspended in 10 ml PBS and exposed to UV light (365 nm) for 1 hour at 4¤ C. The cells were then re-suspended in 10 mM Tris pH 8, 0.25 % lithium diiodosalicylic acid (LIS), 2% Triton x100, sonicated, the cell debris were spun down, and the supernatant ultracentrifuged (200,000 × g for 45 min 4¤ C) the pellet was re-suspended in PBS pH=6.5, 5 mM EDTA, 2% n-octyl-β-D-glucopyranoside (β-OG), ultracentrifuged again – and the colicin with its bound/crosslinked proteins was extracted by EDTA resistant Ni-beads cOmplete (Merk). The extracted proteins were run on 12%-SDS page gels, GFP fluorescent bands of adequate size were analyzed by LC MS/MS for crosslinking mapping.

### LC MS/MS crosslinking analysis

Peptides were separated on an EASY-nLC 1000 ultra-high-performance liquid chromatography (UHPLC) system (Proxeon) and electrosprayed directly into a Q Exactive mass spectrometer (Thermo Fisher). Peptides were trapped on a C18 PepMap100 pre-column (300 μm inner diameter × 5 mm, 100 Å pore size, Thermo Fisher) using solvent A [0.1% (v/v) formic acid in water] at 500 bar and then separated on an in-house packed analytical column (50 cm × 75 μm inner diameter packed with ReproSil-Pur 120 C18-AQ, 1.9 μm, 120 Å pore size, Dr. Maisch GmbH) with a linear gradient from 10% to 55% (v/v) solvent B [0.1% (v/v) formic acid in ACN] in 45 min at 200 nL/min. Full scan MS spectra were acquired in the Orbitrap (scan range 350-2000 m/z, resolution 70,000, Automatic Gain Control target 3e6, maximum injection time 100 ms). After the MS scans, the 10 most intense peaks were selected for Higher-energy collisional dissociation (HCD) fragmentation at 30% of normalized collision energy. HCD spectra were also acquired in the Orbitrap (resolution 17,500, Automatic Gain Control target 5e4, maximum injection time 120 ms) with first fixed mass at 100 m/z. Charge states 1+ and 2+ were excluded from HCD fragmentation. MS data searched using the pLink software^51^. The database contained the target proteins and common contaminants. Search parameters were as follows: maximum number of missed cleavages = 2, fixed modification = carbamidomethyl-Cys, variable modification 1 = Oxidation-Met, variable modification 2 = Glu to pyro-Glu. Crosslinking from D to K, S, T or N-terminus was considered. Data were initially filtered to a False-discovery rate (FDR) of 1%. Crosslinks were further filtered/inspected with specific emphasis on fragmentation patterns.

### Structural modeling

Note: EC (Encounter Complex), SC (Stable Complex)

### Computational modeling of the FepA-ColB interaction

#### Structure preparation

The crystal structures of ColB (1RH1^29^) and FepA (1FEP^20^) were used as starting templates for the computational modeling. Because the crystal structures were missing key loops needed to effectively propagate backbone motions, we added these loops (residues 31-44 on ColB, 323-335 and 384-40 on FepA) using SWISS MODELLER^52^. To eliminate energetically unfavorable side-chain or backbone clashes, we then relaxed the structures using constraints to the native crystal coordinates using RosettaRelax^53^.

### Stage 1: Modeling the semi-rigid encounter complex (EC)

We determined putative local binding conformations by first performing rigid-body global docking using Rosetta’s ReplicaDock2 protocol (built upon prior work on temperature and Hamiltonian replica exchange Monte Carlo approaches^54,55^) and clustering the lowest energy docked structures. Then starting from each low-energy structure, we refine the structures in a local binding region by using our RosettaDock4.0^56^ protocol that adaptively swaps receptor and ligand conformations from a pre-generated ensemble of structures. We diversify the backbone conformations in the ensemble by using: (1) ReplicaDock 2.0 (2) Rosetta Relax^53^ and (3) Rosetta Backrub^57^. Local docking generates ~6,000 decoys, which are scored based on their interface energies, defined as the energy difference between the total energy of the complex and the total energy of the monomers in isolation (see Extended data for detail and command-lines).

### Stage 2: Modeling the stable complex (SC) allowing backbone flexibility

To explore the possibility of ColB flexible N-terminal domain (residues 1-55) interacting explicitly with FepA, we used the Rosetta FloppyTail^58^ algorithm, which allows modest sampling of backbone degrees of freedom following a two-stage approach. First, in the low-resolution stage, side-chains are represented by a centroid atom and the backbone conformational space is extensively sampled. Then, in the high-resolution stage, all side-chain atoms are returned to refine the structures. We generated ~5,000 hypothetical decoys starting from the encounter complex obtained in Stage 1 (EC). The 5,000 perturbation cycles and 1,000 refinement cycles were used for each decoy. To direct the MC sampling of the FloppyTail algorithm toward possible interacting regions, atom-pair constraints based on the experimental (*in vitro)* crosslinking residues guided the search. These constraints were calculated based on a “harmonic” potential with a mean of 6 Å and a standard deviation set to 0.25 Å between the Cα atoms of the candidate residues. Each output decoy was further relaxed to remove unfavorable clashes, and the 100 top-scoring models were then docked using RosettaDock4.0^56^ using a fixed backbone. Translational and rotational moves were performed on the top models to generate ~5,000 docking decoys. To confirm the feasibility of these decoys, we evaluated the interface energies and compared the energy landscape of decoys in Stage 2 with the prior decoys obtained in Stage 1 (Fig. 1c).

### Stage 3: Prediction of the translocation pathway applying *in vivo* crosslinking data

Following the partial unfolding of the plug domain in the related TonB-BtuB system^42^, we allowed backbone movement in the FepA 75-residue half-plug domain (residues 1-75) and the ColB flexible N-terminal domain (residues 1-43). Since simulating the dynamic unfolding of FepA half-plug with simultaneous translocation of the ColB via the barrel protein would be intensely demanding computationally, we instead create models to represent three steps along the dynamic pathway of the unfolding-translocation process. A figure showing the workflow with intermediate snapshots and complete details of each phase of our three-part model creation are given in the Extended data computational methods. Briefly, to create each structure along the pathway, we (1) displace the FepA half-plug (residues 1-75) using Rosetta FloppyTail to pull the terminus out by 4, 8, and 12 Å, respectively, to begin making each of the three structural steps in the pathway; (2) translocate the ColB N-terminal domain (residues 1-43) using both *in vitro* and *in vivo* crosslinking constraints with Rosetta FloppyTail; and (3) refine both FepA and ColB conformation and rigid-body displacement using RosettaDock with a flexible FepA half-plug and ColB N-terminal domain. During stages (1) and (2), backbone motions in FloppyTail are propagated toward the closest terminus, but in stage (3), ColB backbone perturbations during docking are propagated back toward the bulk of ColB to facilitate it finding the optimal rigid-body displacement while the N-terminal domain is translocating. Finally, we calculate interface scores to reveal the favorability relative to conformations of other models presented in this paper along the hypothesized unfolding-translocation pathway (Extended data Fig. 7).

## AUTHOR INFORMATION

### Funding Sources

The experimental work was funded by a BBSRC grant BB/P009948/1. AH and JJG were funded by the U.S. National Institutes of Health grant R01-GM078221.

## Notes

### Competing Interest Statement

The authors have declared no competing interest.

### Summary of Updates

Rearrangement of figure and text order

