## Supplementary information for "Colicin-mediated transport of DNA through the iron transporter FepA"

Extended data

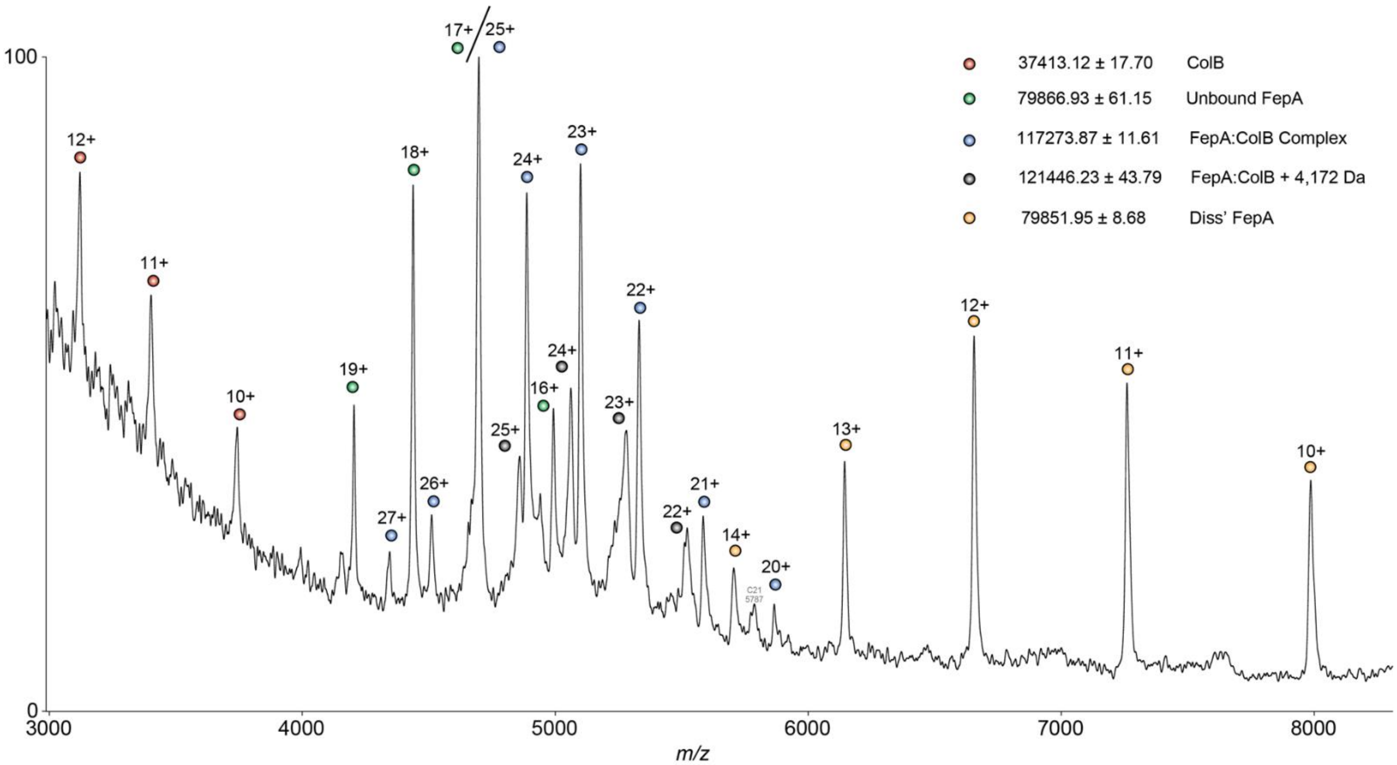


**Extended data Fig. 1| ColB-FepA stoichiometric complex ratio as observed by native state mass-spectrometry.** A clear charge state distributions corresponding to unbound FepA and ColB are observed, as well as a 1:1 non-covalent complex composed of one copy of each protein. Charge reduced species of FepA is also present at higher m/z and indicative of a gas-phase induced dissociation. Also observed is a low abundance charge state distribution which corresponds to the 1:1 / FepA:ColB complex with a discrete mass increase of approximately 4,172 Da. This may correspond to the binding of a single LPS molecule often observed with membrane proteins from the OM, but no further experiments were conducted to further identify the adduct of this low abundance species.

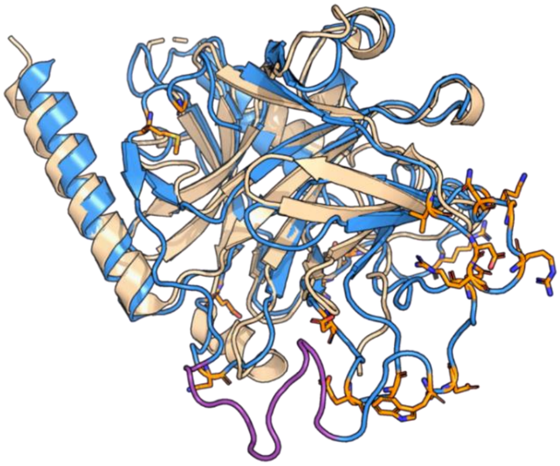

**Extended data Fig. 2| Structural alignment of ColB RT PDB 1RH1 (blue) and ColE7 T domain PDB 2AXC (wheat) and positions of *pBPA* incorporation (orange sticks)**. RMSD 2.384 as calculated by PyMol. Both ColB RT and ColE7 T share a similar pyosin_S fold, yet ColB RT is the only one forming a complex with FepA. *pBPA* has been incorporated mainly in ColB unique surface loops, to examine their role in FepA binding.

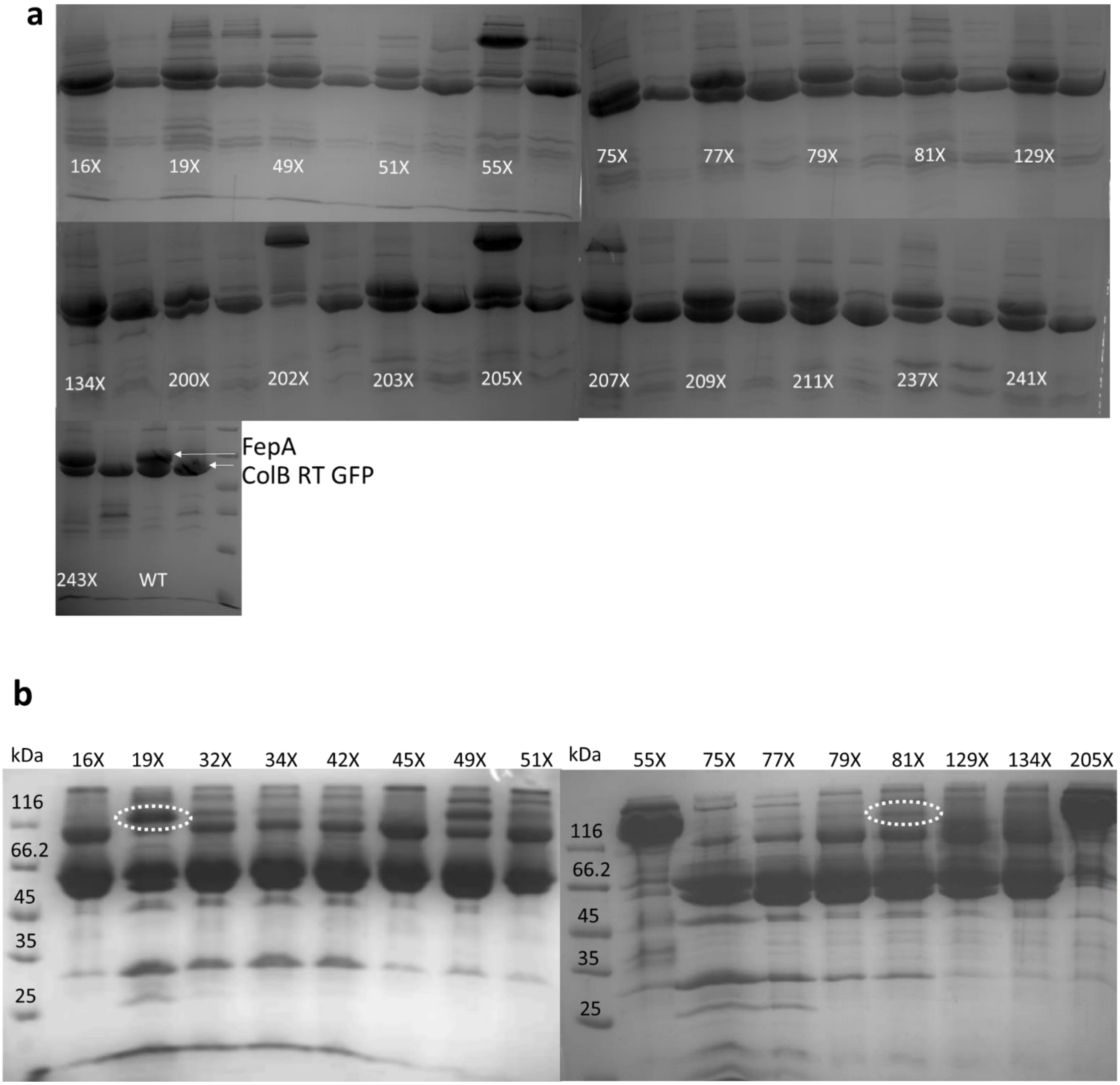
a
16X
19X
49X
51X
55X
75X
77X
79X
81X
129X
134X
200X
202X
203X
205X
207X
209X
211X
237X
241X
243X
WT
FepA
ColB RT GFP
b
kDa
16X
19X
32X
34X
42X
45X
49X
51X
116
66.2
45
35
25
kDa
55X
75X
77X
79X
81X
129X
134X
205X
116
66.2
45
35
25

**c**

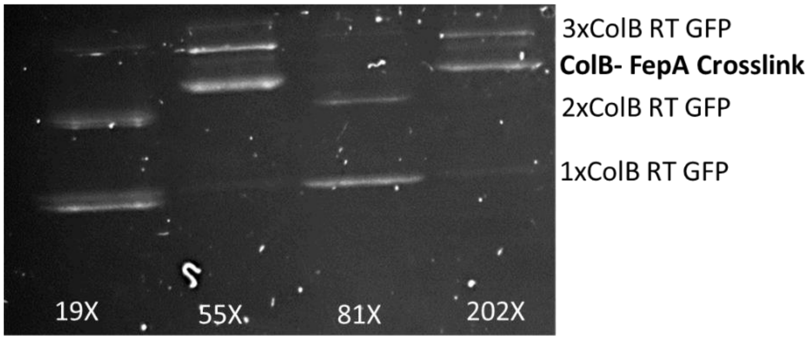


**Extended data Fig. 3| ColB RT *pBPA* GFP crosslinks to FepA. a.** 12% SDS-gel pages emphasizing ColB RT GFP-FepA crosslinking experiments *in vitro*. Crosslinks identified as significant size shifted bends as observed for residues 55, 202 and 205. A self-crosslinking control (with no FepA presence) was run to the right of each lane. **b.** ColB RT GFP-FepA crosslinking experiments performed *in vivo*. **c.** *In vivo* crosslinking experiments using TonB knockout cells (GFP fluorescence image).

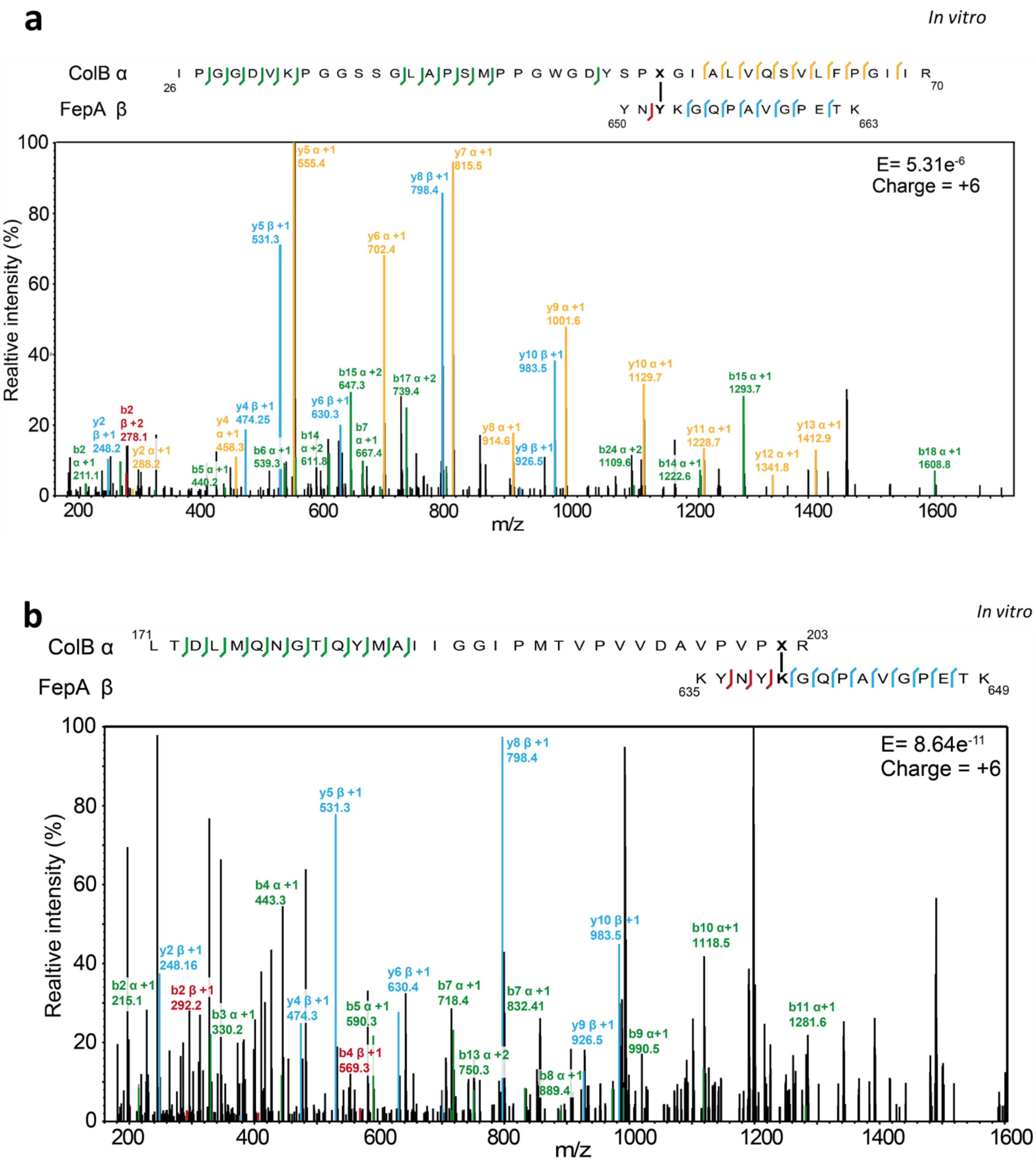

a
In vitro
ColB α
26 I P G G D V K P G G S S G L A P S M P P G W G D Y S P X G I A L V Q S V L F P G I I R 70
FepA β
650 Y N Y K G Q P A V G P E T K 663
E= 5.31e-6
Charge = +6
Realtive intensity (%)
m/z
b
In vitro
ColB α
171 L T D L M Q N G T Q Y M A I I G G I P M T V P V V D A V P V P X R 203
FepA β
635 K Y N Y K G Q P A V G P E T K 649
E= 8.64e-11
Charge = +6
Realtive intensity (%)
m/z

**c** *In vitro*

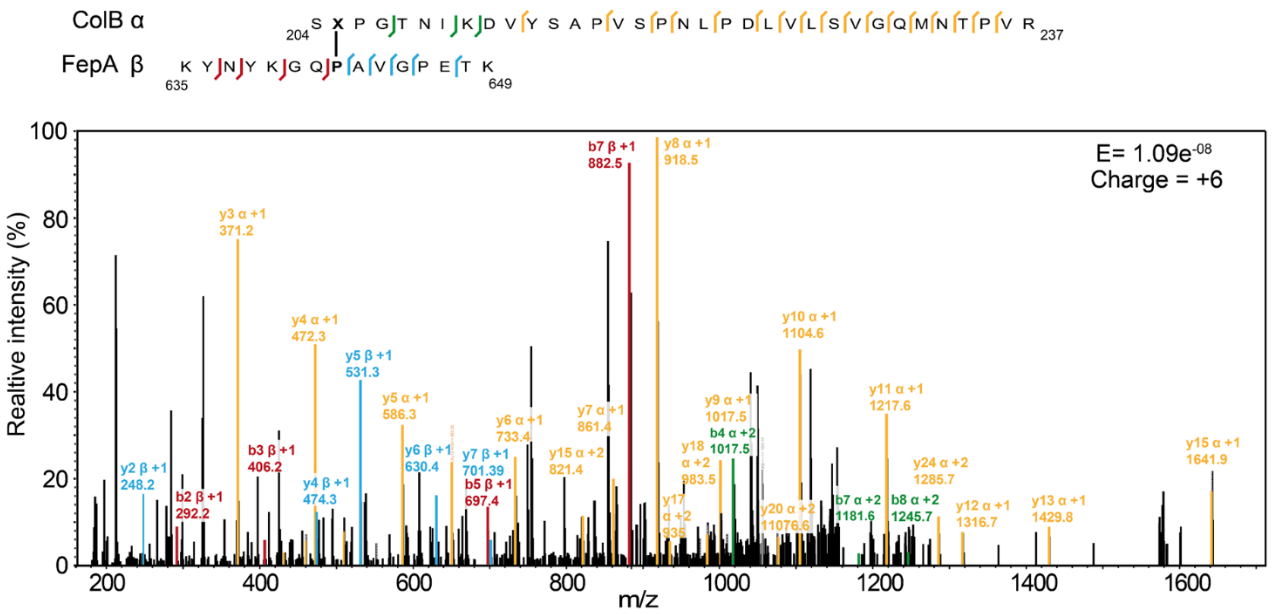


**d** *In vivo*

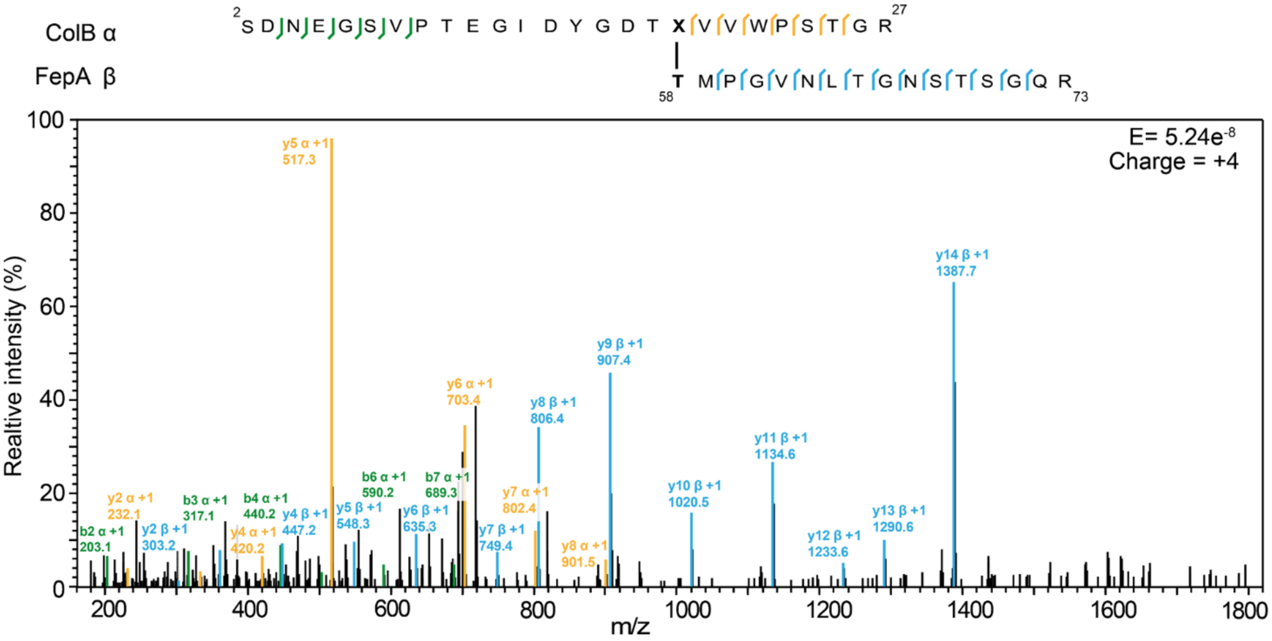

**e**

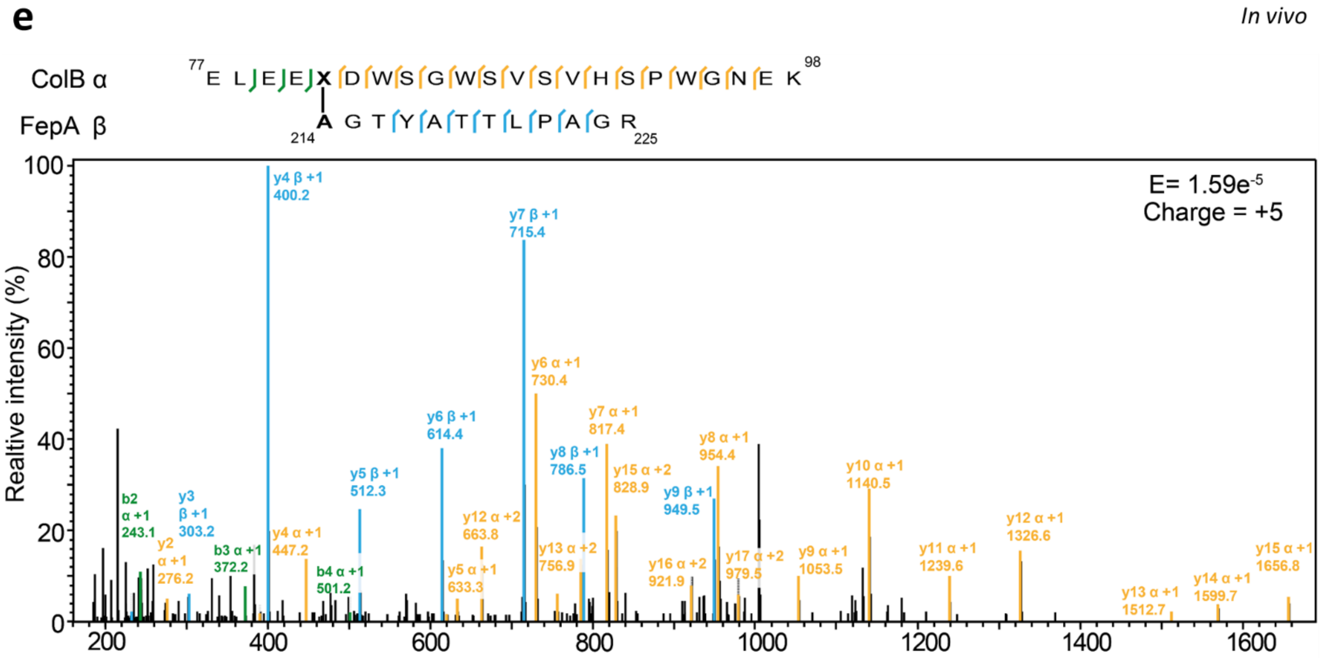


**Extended data Fig. 4| LC MS-MS analysis of the ColB – FepA crosslinks (shown in Extended data Fig. 3) a**. LC MS-MS spectra of ColB RT 55X GFP *in vitro* identified crosslink indicating association with FepA 652Y. **b**. LC MS-MS spectra of ColB RT 202X GFP *in vitro* identified crosslink indicating association with FepA 639K. **c**. LC MS-MS spectra of ColB RT 205X GFP *in vitro* identified crosslink indicating association with FepA 642P. **d**. LC MS-MS spectra of ColB RT 19X GFP *in vivo* identified crosslink indicating association with FepA 58T. **e**. LC MS-MS spectra of ColB RT 81X GFP *in vivo* identified crosslink indicating association with FepA 214A.

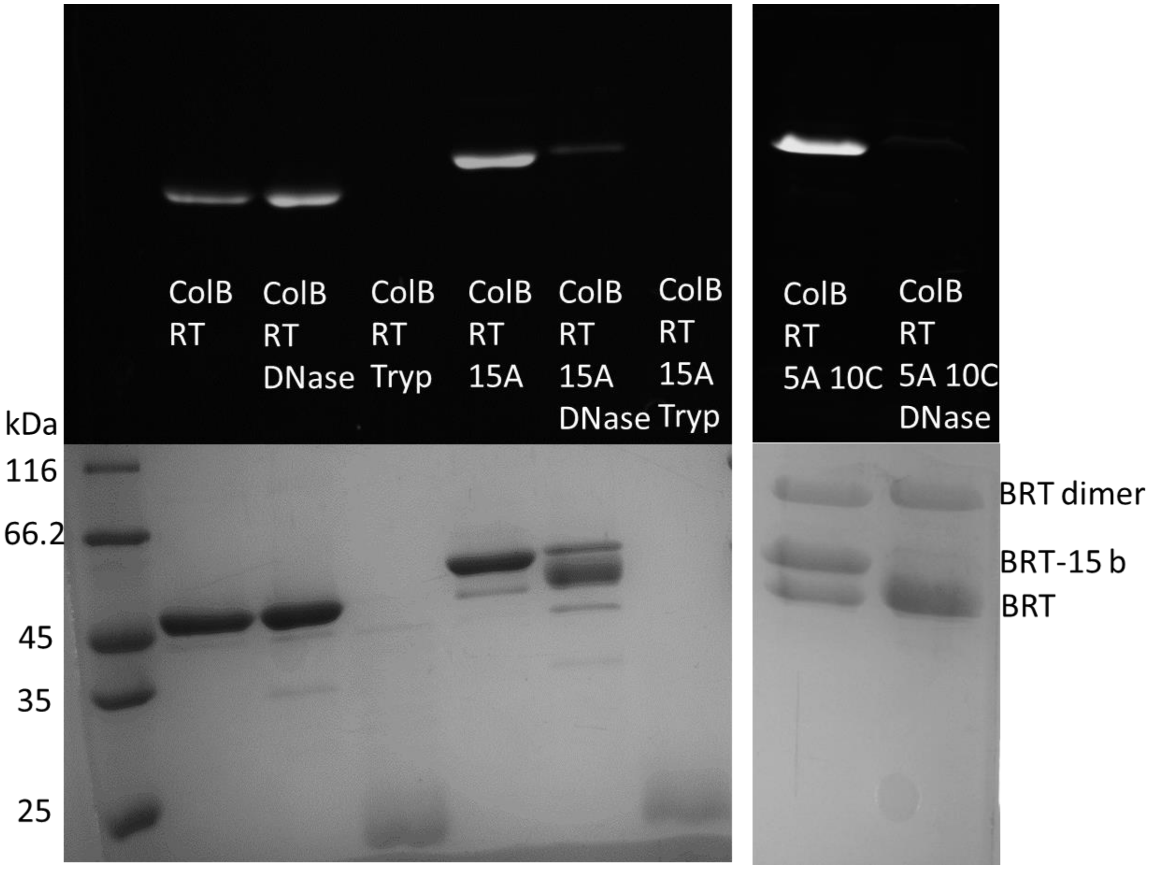


**Extended data Fig. 5| ColB RT conjugation to a fluorescent 15 b ssDNA construct.** ColB RT has either been attached directly to Alexa 488 (left) or via a 15 adenine single-strain DNA construct (center), or via a 5 adenine 10 cytosine ssDNA (right) . Upper panel images Alexa 488 fluorescence, lower panel – coomassie blue protein stain. BRT: ColB RT – Alexa 488, BRT-15 b: ColB RT conjugated to 15 b ssDNA, BRT dimer: non-labeled ColB RT dimers mediated through the introduced C-terminal Cys.

a

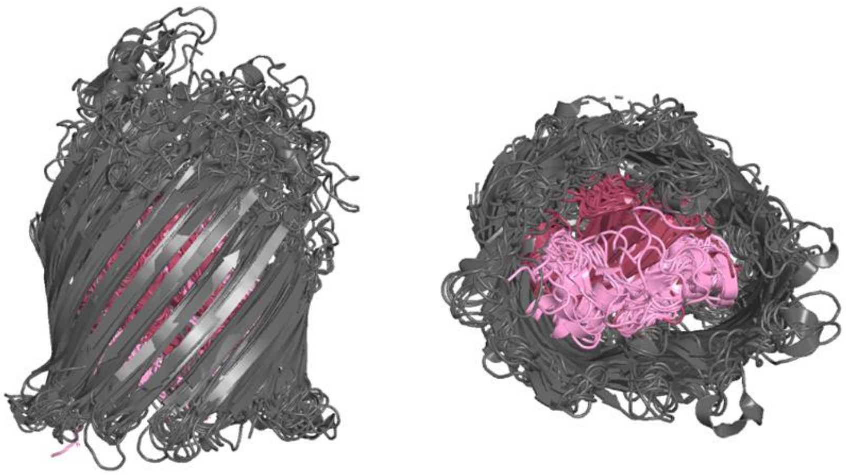

| PDB | RMSD | Protein | Description | Origin |
|---|---|---|---|---|
| 1FEP | 0 | FepA | Siderophore transporter | Escherichia coli |
| 5MZS | 1.23 | PfepA | Siderophore transporter | Pseudomonas aeruginosa |
| 5FR8 | 1.96 | PirA | Siderophore transporter | Acinetobacter menangities |
| 1QJQ | 2.57 | FhuA | Ferric hydroxamate receptor | Escherichia coli |
| 4EPA | 3.14 | FyuA | Ferric yersiniabactin uptake receptor | Yersinia pestis |
| 1NQH | 3.19 | BtuB | B12 Transporter | Escherichia coli |
| 1PO0 | 3.26 | FecA | Ferric citrate transporter in complex with iron-free citrate | Escherichia coli |
| 2HDI | 3.41 | Cir | Colicin I receptor Cir in complex with RBD of Colicin Ia | Escherichia coli |
| 3FHH | 3.41 | ShuA | Heme/Hemoglobin outer membrane transporter | Shigella dysenteriae |
| 4AIP | 3.48 | FrpB | Iron transporter | Neisseria meningitdis |
| 4RDT | 3.62 | ZnuD | Zn-transporter | Neisseria meningitdis |
| 1XKW | 3.85 | FptA | Pyochelin outer membrane receptor | Pseudomonas aeruginosa |
| 2IAH | 3.93 | FpvA | Ferripyoverdine receptor bound to substrate | Pseudomonas aeruginosa |

b

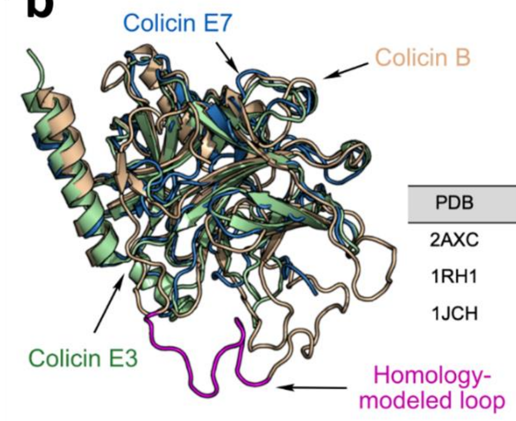


| PDB | RMSD | Protein | Description | Origin |
|---|---|---|---|---|
| 2AXC | 0 | Colicin E7 | N'-terminal translocation domain | Escherichia coli |
| 1RH1 | 2.38 | Colicin B | N'-terminal translocation domain | Escherichia coli |
| 1JCH | 9.42 | Colicin E3 | N'-terminal translocation domain | Escherichia coli |

**Extended data Fig. 6| FepA and ColB RT are representatives of common protein folds. a.** Structural alignment of FepA and 12 additional 22-stranded β-barrel OM bacterial proteins identified by the MADOKA server summarized at the presented table. Side view of the alignment (left) bottom view (right) N-terminal half plug domain in pink, C-terminal half plug domain in hot-pink. **b.** Structural alignment of ColB (blue) with ColE7 (gray) and ColE3 (green) N-terminal translocation domains identified by Pfam as pyocin_S domain superfamily. Table summarizing alignment details at the right hand side.

**Extended data computational Methods**

**PDB Curation and Minimization**

We acquire the crystal structures from the PDB (Colicin B, PDB ID: 1RH1 and enterobactin siderophore receptor FepA, 1FEP). We remove additional water and heteroatoms and clean the PDB structure of all the HETATM and non-canonical amino acids.

```
$ grep ATOM 1fep.pdb > 1fep_clean.pdb
$ grep ATOM 1rh1.pdb > 1rh1_clean.pdb
```

To construct the missing residues, we pass the amino-acid FASTA sequence of the monomer chains with their respectively template PDB structures on SWISS MODELLER, and build models for computational modeling. Before initiating the docking pipeline, we alleviate atomic clashes and optimize the structure with Rosetta Relax.

```
$ relax.linuxgccrelease
      -in:file:s <PDB>
      -relax:thorough
```

**Global Docking**

To determine putative local binding sites, we perform global docking with the Rosetta ReplicaDock2 protocol. The ReplicaDock protocol performs temperature replica exchange MC simulations that allows multiple replicas to communicate with each other and in turn improve the sampling of the energy landscape. We perform global docking in a low-resolution stage while restricting motion to only rigid body moves to better explore the protein energy landscape. To perform global docking, we initiate 8 trajectories of the docking simulation, each trajectory spanning over three temperature replicas run for 2 x $10^6$ MC steps. Inverse temperatures for the replicas are set to $\beta$, of $1.5^{-1}$ $kcal^{-1}$.mol, $3^{-1}$ $kcal^{-1}$.mol and $5^{-1}$ $kcal^{-1}$.mol, and replica exchange swaps are performed every 1000 MC steps. The evaluation is based on interface energies calculated with a six-dimensional, residue-pair transform Motif Dock Score.

### Ensemble Generation methods

A pre-generated ensemble effectively determines the ability of the conformer-selection approaches to select the most promising backbone conformations that can form a thermodynamically feasible complex structure. We use three methods to sample diverse backbone structures.

### ReplicaDock 2.0 (Induced-fit moves)

We built and benchmarked a new method incorporating temperature and Hamiltonian REMC along with induced-fit motions in docking. In this local docking approach, we capture backbone motions of putative interface residues on-the-fly while docking. We perform the search on 8 trajectories all initiated at randomly oriented local binding sites, and each trajectory spans over three replicas run for $2 \times 10^5$ MC steps. We set the temperatures, $1/\beta$, of $1.5\,kcal.mol^{-1}$, $3\,kcal.mol^{-1}$ and $5\,kcal.mol^{-1}$ for the low, medium and high temperatures respectively and replica exchange swaps are attempted every 1000 MC steps. We perform an all-atom refinement over the generated models and the top scoring 50 decoys are seeded into the ensemble. More details on how to utilize the protocol will be incorporated in our future work.

### Relax

We utilize the Rosetta FastRelax protocol to sample backbone conformations of the monomers in isolation. The relax protocol is an equilibration protocol that performs side-chain packing, all-atom refinement and optimization in torsional space. For each monomer, we generate 25 decoys to seed the ensemble.

```
$ relax.linuxgccrelease
      -in:file:s <PDB>
      -relax:fast
```

### Backrub

To perform backbone flexing of the protein fragments, we use the Rosetta Backrub protocol that samples orientations about an axis defined by the pivot atoms i.e. start and end atoms of the fragment. This is performed in isolation for the ligand and receptor chains. We seed the ensemble with 25 backrub outputs.

```
$ backrub.linuxgccrelease
      -in:file:s <PDB>
      -backrub:mc kt 0.6
```

### Docking simulations

Upon generating the ensemble, we follow the pre-packing and docking steps of the RosettaDock 4.0 protocol to perform docking. The details are elaborated in prior work by Marze et al. The command line options for this case are as follows:

### Prepacking

```
$ docking_prepack_protocol.linuxgccrelease
      -in:file:s <PDB>
      -ensemble1 <Receptor Ensemble List>
      -ensemble2 <Ligand Ensemble List>
      -partners A_B
```

**Docking**

```
$ docking_protocol.linuxgccrelease

      -in:file:s <PDB>

      -in:file:native <Reference PDB>
            # this is for estimation of I_rmsd and L_rmsd metrics

      -ensemble1 <Receptor Ensemble List>

      -ensemble2 <Ligand Ensemble List>

      -partners A_B

      -docking_low_res_score motif_dock_score

      -mh:path:scores_BB_BB <Path to the MDS tables directory>

      -mh:score:use_ss1 false
```

**All-atom Docking Refinement**

```
$ docking_protocol.linuxgccrelease

      -in:file:s <PDB>

      -in:file:native <Reference PDB>
            # this is for estimation of I_rmsd and L_rmsd metrics

      -partners A_B

      -docking_local_refine

      -ex1 -ex2aro -rebuild_disulf true

      -detect_disulf true
```

**Rosetta FloppyTail**

We adapt the Rosetta FloppyTail protocol elaborately described in Kleiger et. Al[57], Crawley et al. and Zhang et al. To summarize, FloppyTail samples larger conformational changes by perturbing the backbone dihedral angles via small and/or shear moves in low-resolution and all-atom stages. Each FloppyTail cycle undergoes gradient-based optimization and outputs a 'refined' decoy.

```
$ FloppyTail.linuxgccrelease

      -in:file:s <PDB>

      -ex1 -ex2 -use_input_sc


      # to avoid changing the AA identities during packing

      -packing:repack_only

      -run:min_type dfpmin_armijo_nonmonotone


      # Defines the residues to move

      -in:file:movemap floppy_movemap


      # MC sampling options

      -FloppyTail:shear_on 0.25

      -FloppyTail:publication false

      -FloppyTail:refine_repack_cycles 10

      -FloppyTail:perturb_cycles 5000

      -FloppyTail:refine_cycles 1000
```

The movemap file defines the backbone degrees of freedom that are set for the residues. An example of a sample file that moves the residues (residue numbering follows Rosetta Standard numbering style i.e. they are continuously number from 1 based on their appearance in the PDB file).

```
RESIDUE * CHI     # defaults packing

JUMP * NO         # No movement of subunits with respect to each other

RESIDUE 1 76 BBCHI
```

We have also defined constraints based on the experimental crosslinking data and these constraints effectively direct the sampling of backbone degrees of freedom in relevant search space. We use harmonic constraints and the constraint file is as follows:

```
# Constraint files also follow Rosetta Numbering
AtomPair CA 48A CA 10B HARMONIC 6.0 0.25
AtomPair CA 204A CA 72B HARMONIC 6.0 0.25
AtomPair CA 629A CA 193B HARMONIC 6.0 0.25
```

**Computational Workflow for ColB-FepA complex prediction**

A. Workflow for identifying semi-rigid encounter complex (EC)

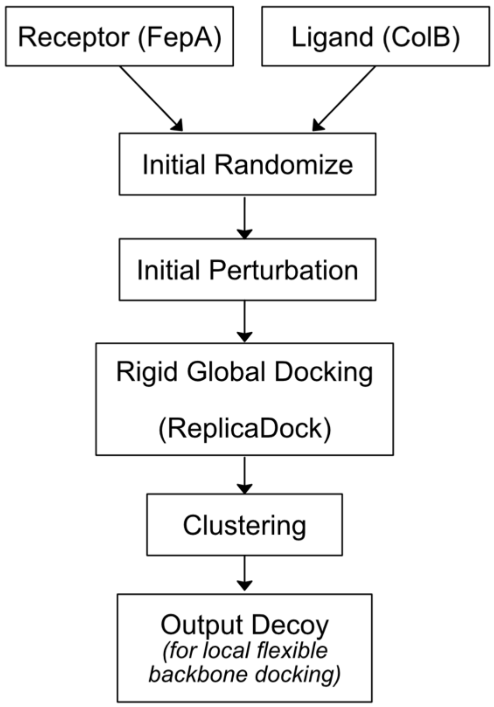

B. Workflow for modeling the flexible encounter complex (SC) between FepA-ColB.

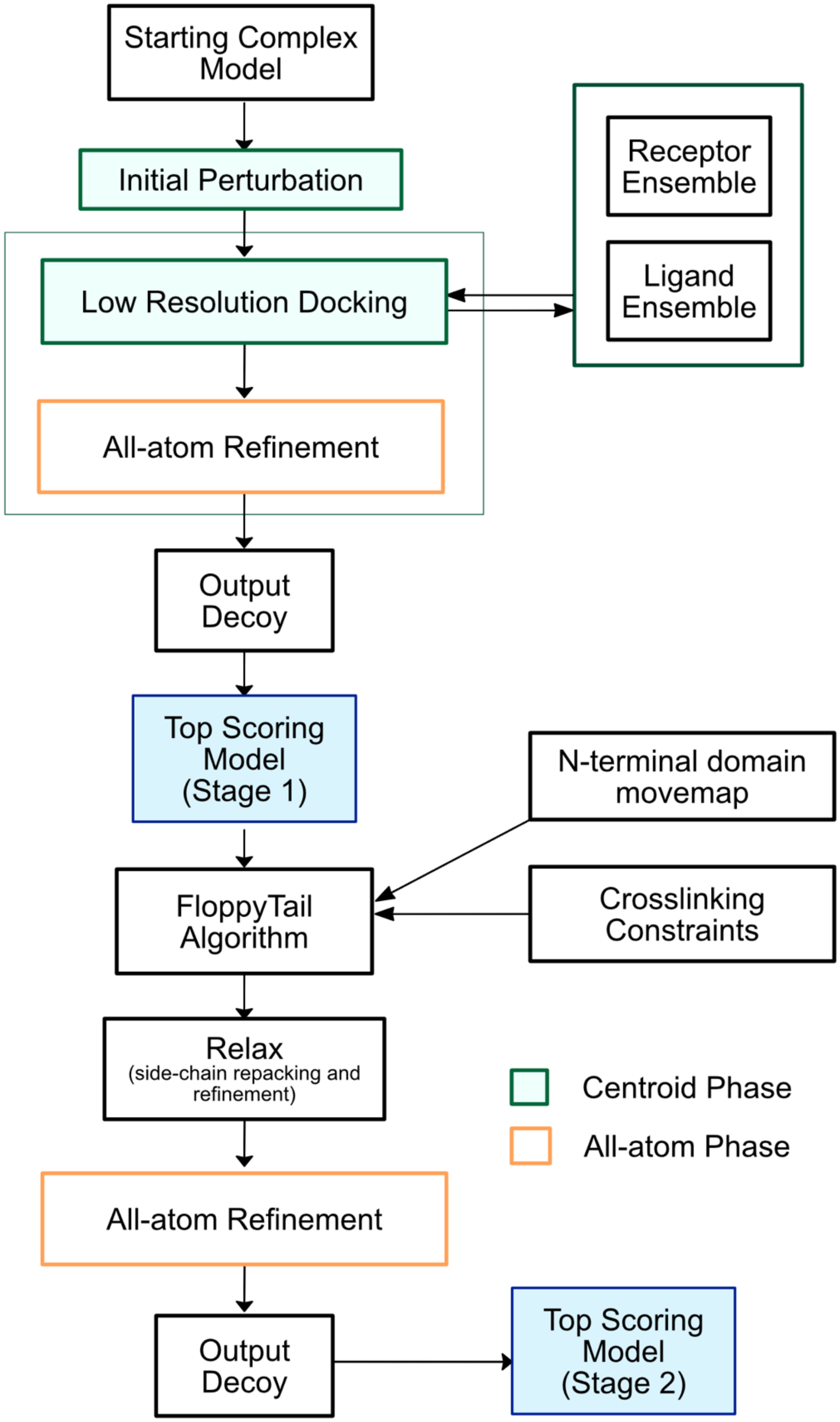

C. Workflow for modeling the complexes in the translocation pathway

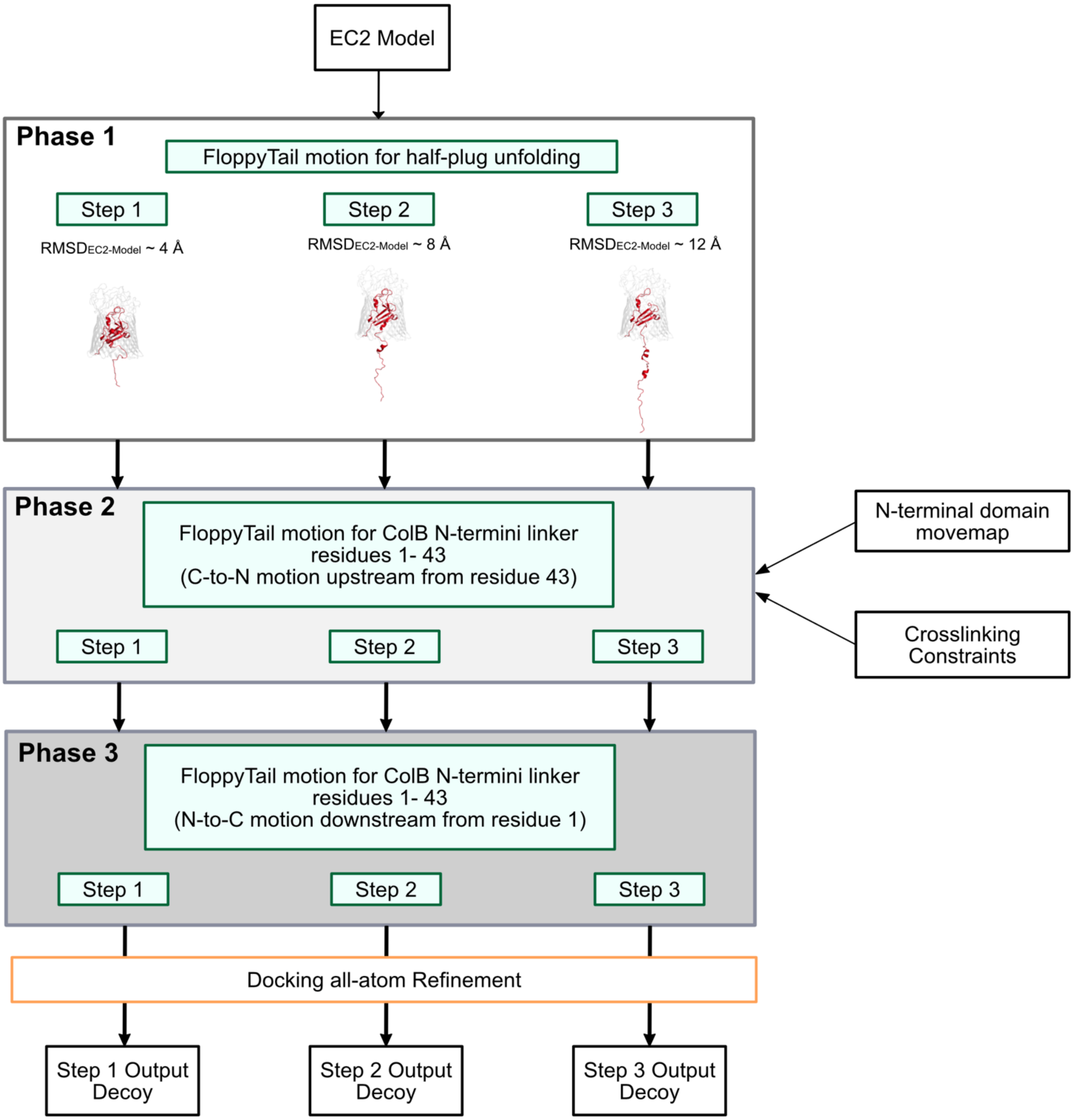

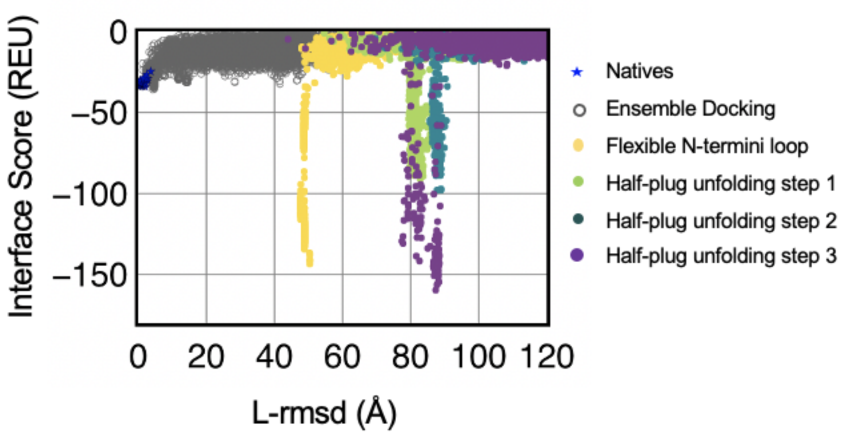


**Extended data Fig. 7| Interface score v/s L-rmsd (Å) for all the decoys generated for the docking simulations.** The global docking minima (blue) are obtained from global docking runs for reference. Stage 1 ensemble docking models (to create EC) are represented in gray and the Stage 2 models (for SC) obtained with FloppyTail are represented in yellow. Stage 3 models involving 3 steps of half-plug unfolding are represented in green (step 1), teal (step 2) and purple (step 3) respectively. As the half-plug is completely unfolded, the interface energies of the colicin B in a partial translocation stage with FepA has a deeper energy well than Stage 2 encounter complex.
